# Rapid and cost-effective development of stable clones for the production of anti-Ebola monoclonal antibodies in HEK293T cells

**DOI:** 10.1101/2020.04.21.054429

**Authors:** Everardo González-González, Iñaki Palestino-Díaz, Felipe López-Pacheco, Alan Roberto Márquez-Ipiña, Itzel Montserrat Lara-Mayorga, Grissel Trujillo-de Santiago, Mario Moisés Alvarez

## Abstract

The Ebola virus (EBOV) disease has caused serious and recurrent epidemics in recent years, resulting in a fatality rate of nearly 50%. The most effective experimental therapy against the EBOV is the use of monoclonal antibodies (mAbs). In this work, we describe the development of HEK293T cells engineered for the transient and stable expression of mAb13C6, a neutralizing anti-EBOV monoclonal antibody. We transfected the HEK293T cells with a tricistronic vector to produce the heavy and the light chain of the antibody 13C6 and intracellular Green Fluorescent Protein (GFP) using Lipofectamine 3000. We then selected the transfected cells using puromycin pressure, dilution cloning, and cloning disks. This integrated strategy generated mAb-producing cells in 7 days with a transient expression of ∼1 mg/L. Stable pools were produced after 4 weeks, with expression levels of ∼0.8 mg/L. Stable clones with expression levels of ∼1.8 mg/L were obtained within 10 weeks. The produced antibodies exhibited the expected functionality; they recognized the GP glycoprotein of the Ebola virus in both ELISA assays and cell binding experiments using HEK293T cells engineered to express the EBOV GP at their membrane surface. By the combined use of GFP and the set of selection techniques here described, we drastically reduced the time from transfection to stable clone generation without resorting to costly equipment. In outbreaks or emergencies, this platform can significantly shorten the development of new biopharmaceuticals and vaccines.

## Introduction

The two major Ebola virus (EBOV) outbreaks occurred since 2013, resulting in 31,737 cases and 13,397 deaths.^1^ The disease caused by the EBOV exhibit a fatality rate of nearly 50%, but values up to 90% were encountered in some regions.^1^ Currently, no approved treatment or vaccine exists for EBOV, so the standard of care recommended by WHO for EBOV patients consists mainly of the use of analgesics, anti-inflammatory drugs, antibiotics, antiemetics, antihypertensives, and anti-epileptic drugs.^2^ Pharmaceutical companies and research groups are continuing their efforts to develop potential preventive and therapeutic anti-EBOV compounds and to run clinical trials.

At present, the most promising alternatives for specific EBOV treatment are monoclonal antibodies (mAbs), and some therapies based on mAbs cocktails (i.e., ZMapp, Remdesivir, MAb114, and REGN-EB3) are currently in clinical trials.^3,4^ The mechanism of action of these cocktails is simple: specific mAbs recognize and bind to the EBOV, thereby interfering with virus replication and inhibiting or neutralizing the infection.^5,6^ The EBOV GP glycoprotein is the main affinity target of anti-EBOV mAbs, and different mAbs bind to different GP regions.^7,8^ The aim of using an antibody mixture is to target the virus at different sites. ^9,10^

The first cocktail used in humans, and the most studied so far is Zmapp, which is composed of three murine chimeric antibodies, 13C6, 2G4, and 4G7.^11^ Studies conducted *in vitro*, in animal models, and in clinical trials have demonstrated the effective action of the mAbs contained in Zmapp against EBOV.^12–15^ The ZMapp mAbs used in clinical trials were produced in tobacco (*Nicotiana benthamiana*) leaves. Plants are promising hosts for mAbs and other recombinant proteins;^7,16^ however, the principal limitation of the use of plants as pharmaceutical biofactories is generating stably transformed plants with high expression levels of the recombinant proteins. For instance, the time required for the generation of a stably transformed tobacco plant is about 20 weeks.^17^

Recently, other hosts have been considered for the experimental production of anti-EBOV antibodies, including cattle,^18^ equines,^19^ and Chinese hamster ovary (CHO) cells.^20^ Among these alternatives, mammalian cells are an obvious and attractive choice for the production of anti-EBOV mAbs, as mammalian cells can produce complex and large glycoproteins and can generate glycosylation patterns that more closely resemble those induced by viral infection in human cells.^21^ The mammalian cell lines used most frequently in the biopharmaceutical industry to produce protein therapeutics are the CHO, NS0, Sp2/0, PER.C6, and HEK293 cell lines. Among these, PER.C6 and HEK293 are of human origin^22^ and therefore offer further advantages for generating human glycosylation patterns.

The HEK293 cell line was established from a primary embryonic human kidney and transformed with human adenovirus type 5 DNA. It is the most common cell line used to produce recombinant proteins, second only to the CHO cell line.^23^ HEK cells have been used as an expression tool for more than 35 years because of their ease of culture and the feasibility of efficient genetic engineering^24^ for producing transient expression clones that often yield enough protein for research purposes and preliminary characterization studies during drug development. However, producing high levels of protein in mammalian cells, is necessary to generate stable cell lines ^25^; and the time to generate stable cells is long, around 6 to 12 months ^26^

Through the years, many strategies have been proposed and experimentally demonstrated to accelerate the process of generation of stable clones capable of producing reasonably good quantities of pharmaceutical compounds for further testing. In this study, we combine different technologies from the genetic engineering fields, including the use of tricistronic vectors and a cloning selection process based on the combined use of dilution cloning and cloning discs, to generate a stable HEK293T cell line engineered to produce the anti-Ebola mAb13C6.

Reports of stable mAb production in HEK293 cell lines are very infrequent in the specialized literature. Therefore, we believe that the information presented here may be of use to cell engineers both in academia and industry. The proposed strategy is simple and reduces complexity and the time required to generate stable cell lines, while negating the need for costly equipment and materials.

## Materials and methods

### Cell culture

The HEK293T cell line (human embryonic kidney cells from ATCC- CRL-3216) was cultured in Dulbecco’s Modified Eagle Medium Medium (DMEM) (ATCC 30-2002) supplemented with 10% (v/v) fetal bovine serum (FBS; Gibco, MA, USA, Cat. No.16000044). Cells were cultured in T25 and T75 flasks with 5 mL and 15 mL of the prepared medium, respectively. Cultures were maintained in an incubator containing 5% CO2 at 37 °C. Cell culture experiments were run for 7 days for transient expression, 20 days for stable pool culture, and 7 days for stable clone culture.

### Cloning expression vector and DNA purification

We designed pCMV-13C6-2A-GFP as a tricistronic vector for the transfection of mammalian cell lines. This vector, which drives the production of the anti-EBOV mAb13C6 and green fluorescent protein (GFP), was derived from the pCMV-13C6 vector synthesized by ATUM (CA, US). We extracted the antibody gene from pCMV-13C6 vector using polymerase chain reaction (PCR) strategies. To this aim, we used Phusion Green Hot Start II High-Fidelity PCR Master Mix (Thermo Scientific, MA, USA; Cat. No. F566L). Ligation was done using the restriction enzyme *Sap*I in the bicistronic vector pCMV-2A- GFP from ATUM (CA, US). This vector is regulated by a CMV promoter to produce intracellular GFP and secreted 13C6 antibody. We used the same strategy to produce the pCMV-GP-RFP vector, using first the vector pCMV-GP synthetized by ATUM (CA, US) and cloning in a target vector that contained the RFP gene. DNA was purified using the Pure Yield Plasmid Midiprep system kit (Promega, WI, US Cat. No. A2492). DNA quality and concentration were determined by measuring the ratio of signals at wavelengths of 260 and 280 nm using a NanoDrop instrument (Thermo Scientific, MA, USA; Cat. No. ND- 2000).

### Cell transfection

Lipofectamine 3000 (Invitrogen, CA, USA. Cat. No L3000015) was used for the transfection of HEK293T cells. At 24 h before transfection, 4 × 10^5^ cells were seeded in 12-well plates in Opti-MEM Reduced Serum Medium (Gibco, MA, USA, Cat. No. 31985070). For each transfection, 1.25 μg of DNA and 125 μL Opti-MEM, 2.5 μL P3000, and 2.5 μL Lipofectamine 3000 were used per well. The mixture of DNA, P3000, and Lipofectamine 3000 was incubated for 15 min to obtain the complex liposome-DNA. The liposomes were then added dropwise to the cells in the plate. Transfection efficacy was determined after 48 h using fluorescence microscopy (Nikon ECLIPSE Ti2, NY, USA). To that aim, we calculated the ratio of the number of cells that expressed green fluorescence from GFP (wavelength of excitation 470/40 and emission 525/50) under fluorescence microscopy as a percentage of the total number of cells observed in bright field.

### Pressure and selection of stable cells

We used a combination of techniques to identify and isolate mAb13C6 high producer cells from our cultures. This combination of strategies consisted of (a) conventional antibiotic pressure using puromycin, (b) the use of dilution cloning protocols, and (c) disk cloning techniques. In order to determine puromycin (Gibco, MA, USA, Cat. No. A1113803) resistance, we seeded cells in 12-well plates and exposed them for 72 h to different puromycin concentrations, ranging between 0.2 and 5 μg mL^−1^.

Stable pools of cells were obtained by selection pressure with puromycin using selection cycles of 0.8 μg mL^−1^ for 3 weeks (including culture expansions). Stable clone cultures were obtained by combining dilution cloning and the use of cloning discs for the selection of monoclonal cells with high production of GFP. Levels of GFP production were monitored by fluorescence microscopy, both to evaluate the success of the puromycin selection and the culture growth.

### Optical microscopy

Transient cultures, stable pools, and clonal cultures of recombinant HEK293T cells engineered to produce mAb13C6 and GP were periodically analyzed by optical microscopy using an Axio Imager M2 microscope (Zeiss, Germany) equipped with Colibri.2 LED illumination and an Apotome.2 system (Zeiss, Germany). Bright-field and fluorescence micrographs were used to document the progression of the expression level of the different cultures through time. Wide-field images (up to 20 cm^2^) were created using a stitching algorithm included as part of the microscope software (Axio-Imager Software, Zeiss, Germany) to facilitate image analysis.

### Quantification of antibody titer

Quantification of antibody production in the supernatant of all our HEK293T cell cultures was done using an ELISA assay specific for human IgG (Bethyl Laboratories, TX, USA, Cat. No. E80-104) and a standard curve in the range of 7.8–500 ng mL^−1^. The antibody concentration was calculated based on this standard curve. Assays were conducted according to the manufacturer’s instructions.

### Affinity assays using GP cells

We also tested the affinity of the produced mAb13C6 to GP-EBOV in cell culture assays. To do this, we engineered HEK293T cells to express GP protein from EBOV at their membrane surface. Briefly, HEK293T were transfected with a bicistronic vector containing information for the trans membrane expression of RFP-GP (red fluorescent protein fused to GP). HEK293T cells expressing GP at their surface were taken to confluence in 12-well plates, supernatant from HEK293T cells producing mAb13C6 was added, and the cells were incubated for 60 m. The supernatant was then discarded and the adhered cells were washed three times with fresh PBS. The GP-13C6 complex was identified by adding a polyclonal human anti-IgG antibody conjugated to FITC (Abcam, Cambridge, USA; Cat. No. ab97224). Cell cultures were observed under fluorescence illumination with an optical microscope (Axio Imager M2 microscope (Zeiss, Germany) using a set of filters (red and green) to reveal the presence of a red signal (for GP alone) or a combined red and green signal (for the GP-mAb13C6 complex).

## Results and discussion

One important bottleneck in the process of massive production of mAbs (or for developing novel mAbs) is the generation of stable clones^25,27^. The time to generate stable cells is long^27^, at approximately 6 to 12 months ^26^. Here, we demonstrate a simple strategy for generating stable cell lines of HEK293T cells capable of producing an anti-EBOV monoclonal antibody (i.e., mAb13C6) in a short time (less than 3 months). MAb13C6 is one of the monoclonal antibodies present in Zmapp, a promising mAb cocktail that was employed to treat a number of EBOV patients under compassionate use provision during the West Africa EBOV epidemics in 2014. Back then, Zmapp was produced by means of transient expression in tobacco leaves, a process that seriously limited large-scale production.

Our strategy is based on the combined use of (a) tricistronic vectors (also coding for GFP expression for monitoring purposes) and (b) puromycin selection (using dilution cloning and cloning discs). Our methods of transfection and selection use common reagents and equipment frequently found in a typical mammalian cell culture laboratory. By this use of inexpensive and widely accessible equipment and reagents, we aim to enable other research groups currently facing constrained development budgets or restrictive timelines.

### Construct design and cell engineering

We generated HEK293T cells capable of producing both 13C6 anti-EBOV GP antibody and GFP under the command of a unique genetic construct. We used our tricistronic vectors as an alternative to produce three different proteins (the two chains of mAb13C6 and Green Fluorescent Protein (GFP)) regulated under a single promoter. The particular strategy used here to achieve tricistronic expression is schematically presented in Figure 1C. The light and heavy chains of mAb13C6 were linked through an IRES sequence. In addition, we included a short linker (i.e., 2A-peptide) between the heavy chain of mAb13C6 and GFP to enable expression of the triad of proteins under the command of a single promoter. Elements such as an internal ribosome entry (IRES) and 2A-peptide between genes provide the ability to translate more than one protein.^28,29^ GFP offered an alternative to visualize the transfection efficiency and expression during the selection process and greatly facilitated the identification of the cells actually producing the protein of interest.^30^

**Figure 1.**
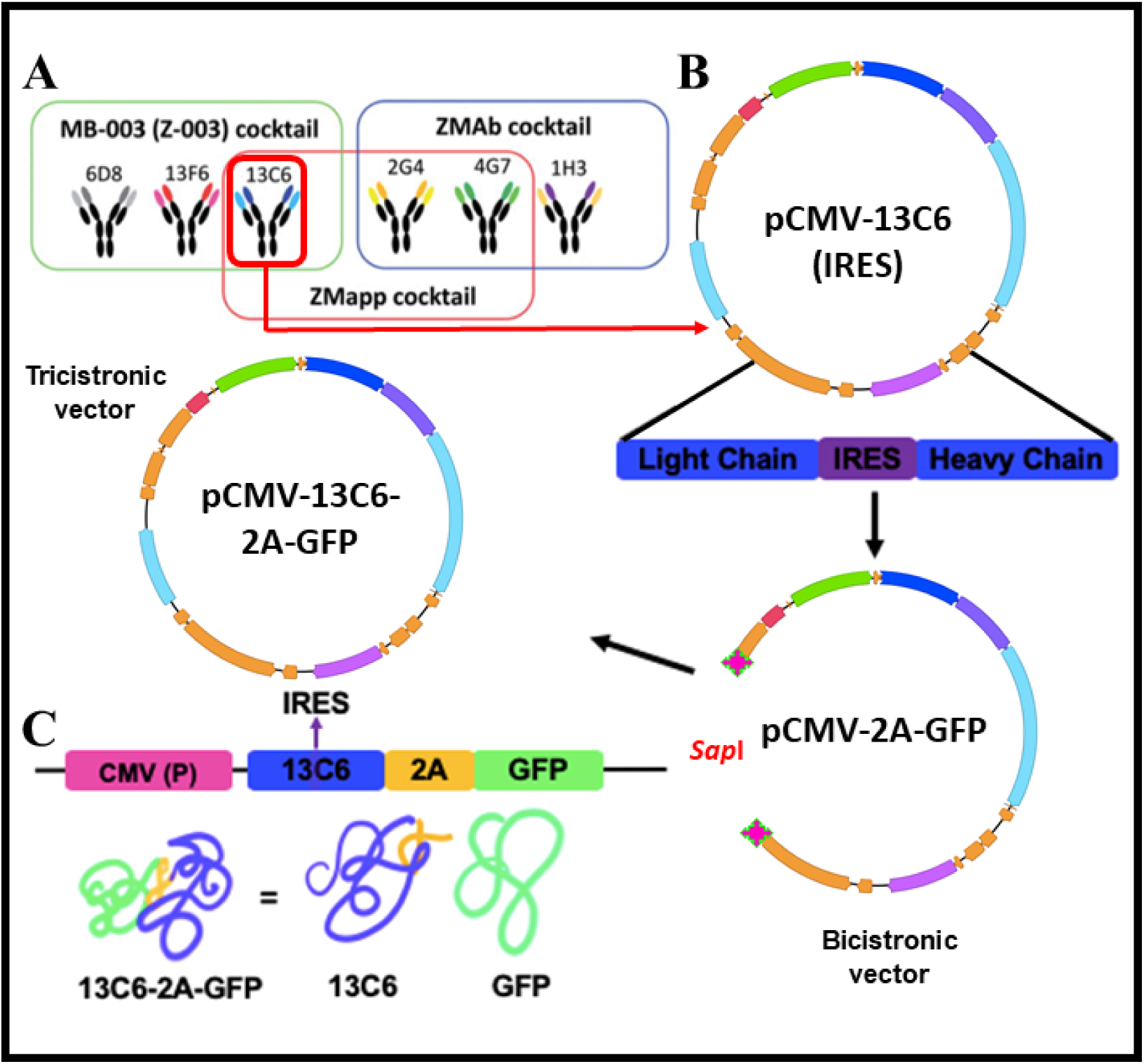
Description of the construction strategy of the tricistronic vector for expression of mAb13C6 and GFP (pCMV-13C6-2A-GFP). **A)** mAb therapeutic cocktails tested against EBOV and the selected antibody for production in this work (mAb13C6). The expression strategy is based in the construction of a tricistronic vector expressing the heavy and light chains of mAb13C6 and GFP as a fluorescent marker of expression. **B)** Cloning strategy to construct the tricistronic expression vector: Extraction of the mAb13C6 gene that includes an IRES sequence from a pCMV-13C6 vector, and for cloning of the IRES-13C6 sequence using the *Sap*I enzyme into a bicistronicvector containing a GFP cassette (pCMV-2A-GFP) **C)** Schematic representation of the final expression vector CMV (promoter), 13C6mAb (expression cassette containing an IRES element), 2A (viral sequence), GFP (Green Fluorescent Protein) and its intended expression: properly folded 13C6mAb and unlinked GFP.

**Figure 2.**
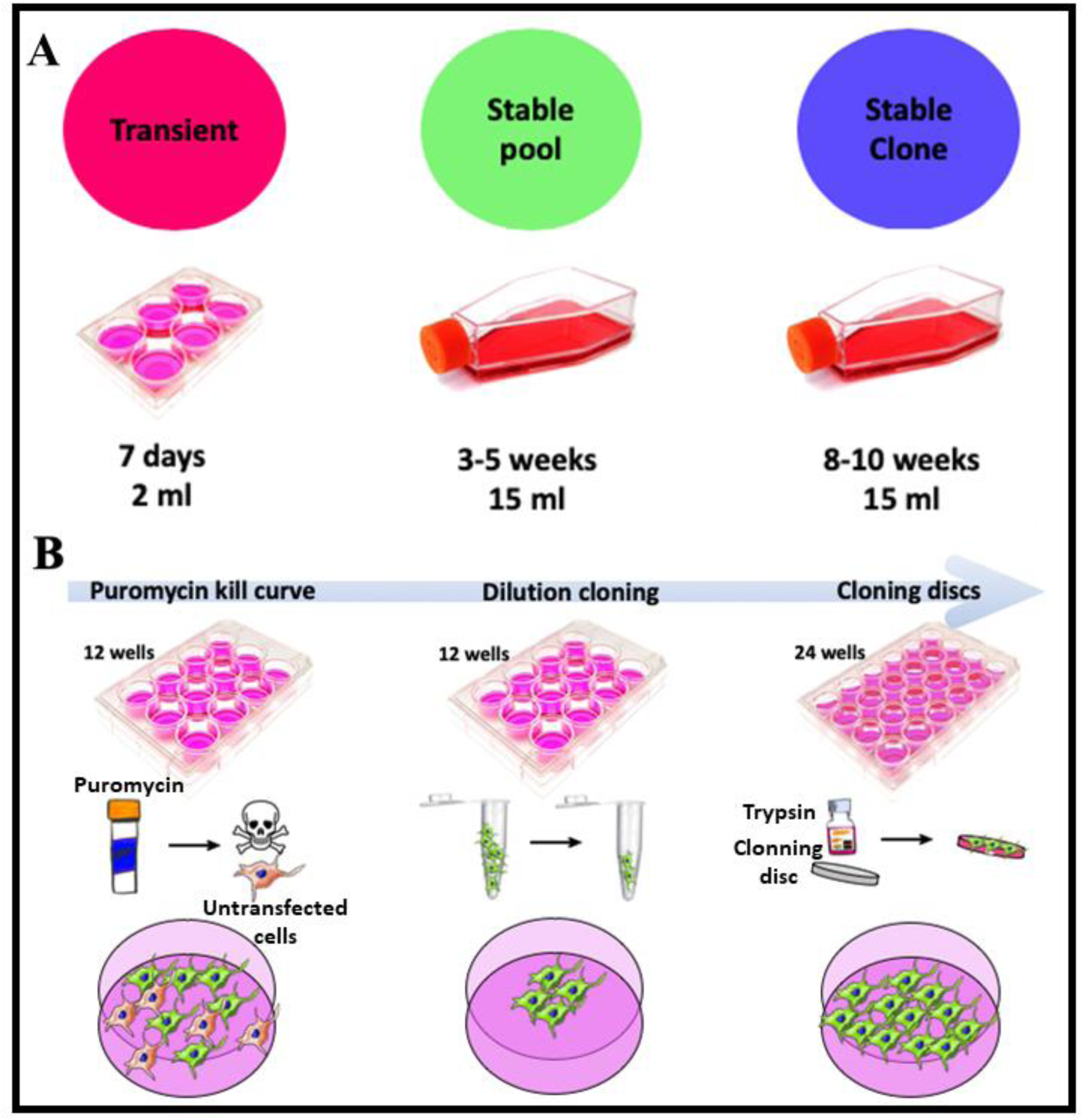
Strategy for generation of transient and stable pools and clones for the expression of mAb13C6. **A)** Production of mAb13C6 in transient, stable pool, and stable clone cultures; time requirements and culture volumes are indicated. **B)** Diagrammatic representation of the pressure and selection process conducted to obtain stable clones for the production of mAb13C6. An optimal puromycin concentration was first determined to select successfully transfected cells. GFP expression was then used for easy monitoring of the growth of successfully transfected cells. Clonal selection was assisted by dilution cloning and the use of cloning disks.

### Transient production of mAb13C6 in HEK293T cells

The generation of mammalian cells capable of transient expression is the first step in the process of developing a recombinant protein of pharmaceutical value. Transient expression enables the production of recombinant protein within only days after transfection, yielding production levels that are often sufficient for characterization and preclinical studies.^27,31–34^ The transfection technique used, in conjunction with the selection of an adequate expression vector (i.e., expression in transient or stable format), are the fundamental factors that determine the success of any strategy aimed at achieving high production of a recombinant protein^27^. Different methods have been reported for transfection of the HEK293 cell line, including the use of polyethylenimine (PEI), electroporation, use of dendrimers, and calcium phosphate precipitation.^23,35–38^

Here, we transfected the CMV-13C6-2A-GFP plasmid using Lipofectamine 3000. Image analysis techniques were used to determine the efficiency of transfection based on microscopy images. The same field of view was first captured under bright field and then under fluorescence illumination. Transfection efficiencies were determined from these pairs of images as the percentage of cells expressing GFP 3 h after transfection. We achieved high efficiencies of transfection (over 80%) in HEK293T cells using this strategy. The conditions and protocols reported here yielded transient cultures 48 h post-transfection.

We determined the concentration of mAb13C6 produced in our transient cultures using ELISA experiments that allowed evaluation of the total concentration of IgGs in the supernatant of transiently expressing HEK293T cells cultured in flasks. For instance, in our experiments, we obtained several milligrams of mAb13C6 at 3 days after transfection (Figure 3A). In our transient cultures, we observed levels of expression of ∼1 mg/L in cultures containing a total of 3 × 10^5^ cells between the 4th and 7th day in a volume of 2 mL (in 6 well-plates) (Figure 3A). The levels of expression that we observed (1.6 ng cell^−1^ day) may be sufficient for most applications related to protein quality assessment. GFP expression was seen 12 days later, which suggests that transient pools can be used to purify small quantities of protein for more than one week.

**Figure 3.**
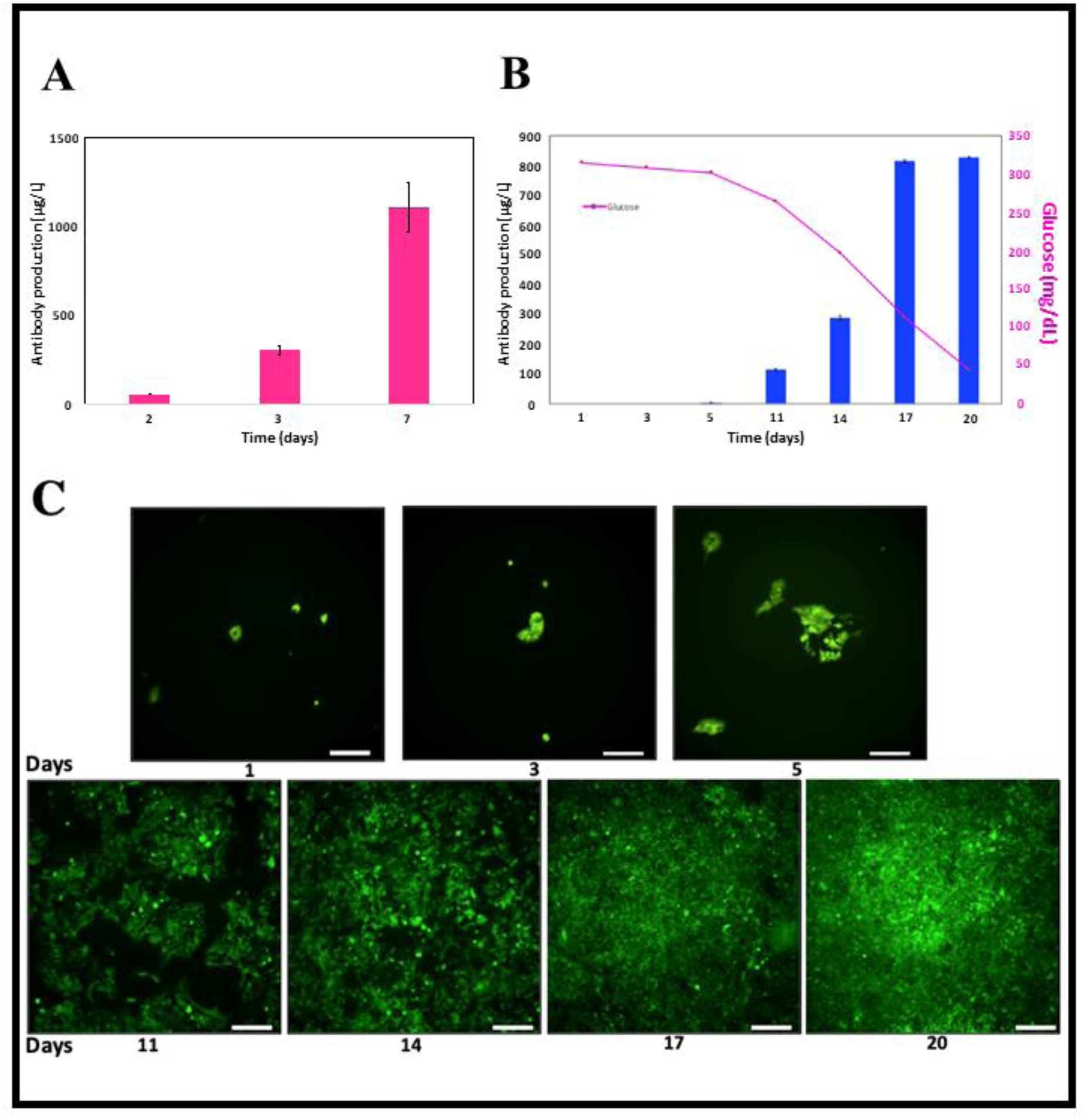
Monnitoring of mAb13C6 production. fluorescence intensity, and cell growth during transient production. **A)** Antibody production in transient culture during a week. **B)** Antibody production and glucose concentration measurements during 20 days in stable pool cultures. **C)** Micrographs showing the GFP fluorescence emission from stable pool cultures over time. Scale bar: = 200 μm.

### Generation of HEK293 cells with stable expression of mAb13C6

While transient expression offers a rapid way to evaluate the potential of a recombinant protein, clinical testing of that protein still requires stable cultures, as does its large-scale industrial production. As stated before, the generation of stable cultures of mammalian cells as producers of monoclonal antibodies is technically cumbersome and lengthy.^27^ Studies related to the development of stable cells typically report a timeline in the range of 6 to 12 months and involve multiple steps of pressure/selection.^27^ New techniques for the generation of stable cell lines have reduced the timeline, but these generally involve the use of costly equipment, such as flow cytometers capable of sorting or automated colony picking (i.e., CloniPix FL).^39–42^

We produced a stable cell pool in less than 5 weeks and stable clones in 10 weeks using the methods described here. We extended the expression and improved antibody production using puromycin to kill low-producer cells and to select for stable pools and clones of high-producer cells (Figure 2A). We first established a puromycin kill curve in the range of concentrations from 0.25–10 μg mL^−1^ (Figure 2B). We obtained the highest fraction of transfected cells and post/treatment viability using a concentration of puromycin between 0.4 and 0.8 μg mL^−1^. Using this pressure strategy, we produced stable pools of mAb13C6 producer cells after 3 to 5 weeks of culture in DMEM medium supplemented with 10% BFS and 0.5 μg mL^−1^ of puromycin.

Stable pools were cultured in T75 flask for 20 days and the concentration profiles of glucose and mAb13C6 were measured (Figure 3B). Continuous monitoring of the glucose concentration allowed us to confirm that cells were metabolically active and that sufficient substrate was still available to sustain viability and mAb13C6 production. The GFP expression was also used as a reporter in our experiments to monitor the viability and overall condition of the cultures. We closely monitored the status of our cultures by visual inspection under fluorescent illumination with an optical microscope (Figure 3C). As before, we determined the concentration of mAb13C6 using ELISA. We observed sustained concentration levels of approximately 0.8 mg of IgG L^−1^ during the last 5 days of stable pool cultures from final cell counts of 1.5 × 10^6^ cells in volumes of 15 mL (Figure 3B). For reference, these values imply specific cell productivities of approximately 1.6 +/- 0.04 pg cell^−1^ day^−1^, or one order of magnitude below those reported for stable and high producer CHO cell clones in typical commercial processes.^43^

We used our stable pool cultures to conduct dilution cloning, and we cultured the clones for an additional week to expand the clones. We then used cloning discs to single out clones, and we grew them in new and independent wells for 1 to 2 weeks before scaling the culture from 24-well to 6-well plates. This strategy allowed us to obtain stable clones after approximately 10 weeks. From the stable pools of mAb13C6 producers, we selected 4 high-expression clones, designated as C66, C95, C96, and C97, for further characterization. Selection was based on the level of fluorescence emitted by the cultures of stable pools, as observed and measured by optical microscopy. As expected, we observed higher production of mAb13C6 in the stable clone cultures than in the stable pools. The selected clones were cultured in T75 flasks using 15 mL of medium for a period of 7 days. The final viable count in these cultures was approximately 1.5 × 10^6^ cells. We determined maximum mAb13C6 titers of 1.8 mg L^−1^ in 3 days. Among the selected clones, the best performers were clones C66 and C95 (Figure 4A).

**Figure 4.**
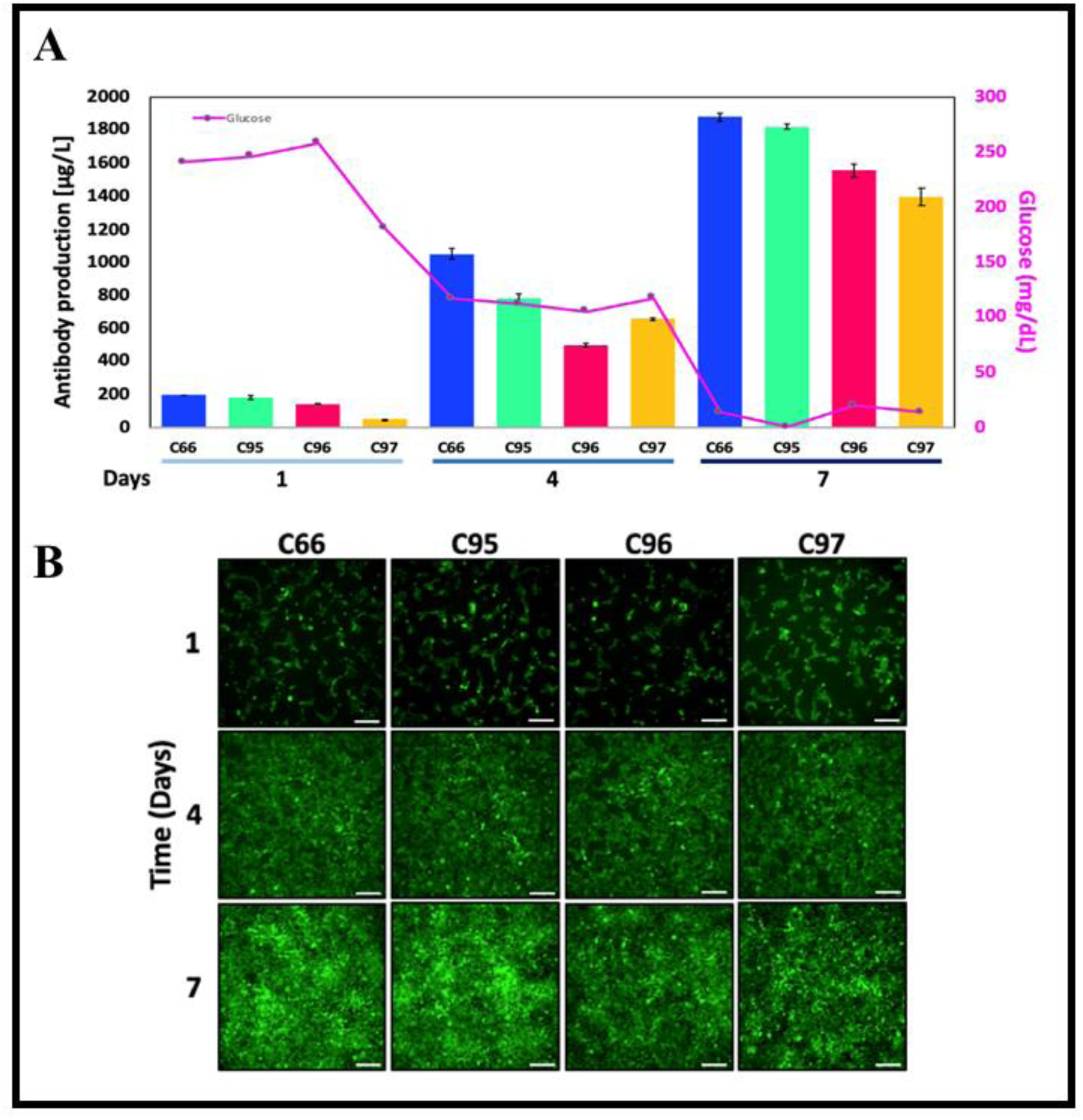
Antibody production in stable clones. **A)** Antibody production and glucose concentration profiles exhibited by the stable clones C66, C95, C96, and C97 during one week of culture. **B)** Micrographs of the stable clones cultured for 1–7 days. Scale bar: 200 μm.

Figure 4C shows mAb13C6 titers and glucose concentrations at different time points during the culture of each of the four stable clones that we characterized. We observed a clear correlation between glucose consumption and mAb13C6 titer as time progressed. As expected, glucose depletion was faster in the high-producer clones than in the stable pool cultures. The stable pool cultures depleted glucose from the culture medium in about 20 days, whereas the high-producer clones consumed all the glucose in only 7 days. As before, GFP expression was used as a reporter for monitoring the cultures and guiding the selection process. Micrographs of the culture progression with time are shown in Figure 4B. The clone cultures became progressively confluent during 7 days of culture without appreciable loss of GFP expression.

In our experiments, we measured the average mAb13C6 concentration values of approximately 1.6 mg L^−1^ among the four selected clones. The mAb titers reported for production in mammalian cells range from micrograms to milligrams per liter.^39,44–46^ The specific cell productivities observed in our four stable clone cultures (5.33 +/- 0.7 pg cell^−1^ day^−1^) were above the range of values obtained in previous studies that used similar or more complex cell engineering strategies on HEK cells. ^35,47,48^

These titer, productivity, and specific productivity values are competitive for experimental cultures but still distant (although in the same order of magnitude) from those expected for commercial scale production (approximately 20–40 pg cell^−1^ day^−1^).^43^

We anticipate that the application of standard optimization protocols (i.e., fed-batch feeding protocols) could enhance productivity by at least one order of magnitude.^49,50^ Further enhancement is possible by translating to suspended culture in instrumented bioreactors^48,51^ or culturing by continuous perfusion in cell adherent culture systems.

### Characterization of the biological functionality of mAb13C6: In vitro testing

We ran PCR to evaluate the presence of the mAb13C6 gene in transient pools, stable pools, and stable clones. We designed specific primers to identify the product of amplification (amplicon) of 3 kb corresponding to the gene that encodes for mAb13C6 expression, and we found the gene associated with the production of mAb13C6 in all our cultures (Figure 5A). We also conducted western blots to visualize the presence of the full antibody. The typical and expected antibody signature profile, namely two bands observed at 50 kDa and 25 kDa, was observed for all our cultures (Figure 5A).

**Figure 5.**
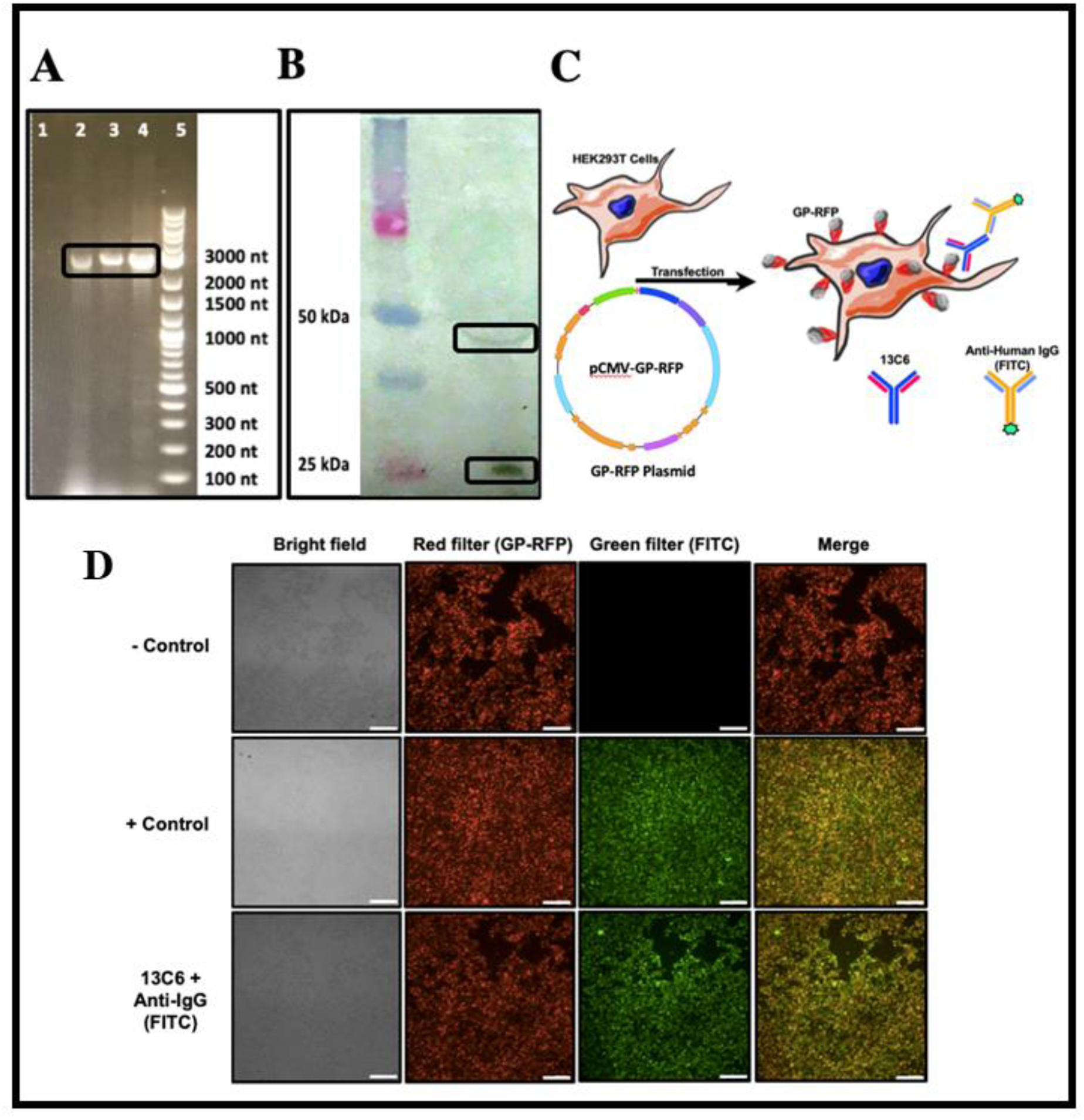
Characterization of the anti-EBOV mAb13C6. **A**) Detection of the gene encoding for the production of mAb13C6 in the different HEK293T cultures: negative control (lane 1); transient cultures (lane 2), stable pools (lane 3), stable clone C95 (lane 4), and reference molecular weight ladder (lane 5). **B)** Western blot showing bands that correspond to the molecular weight of the heavy and light chains of mAb13C6. Proteins were marked with a human anti-IgG antibody. **C)** Schematic representation of the interaction between HEK293T cells expressing GP-RFP at their surface and mAb13C6 marked by a human anti-IgG conjugated with FITC. **D)** Micrographs of HEK293T cells with GP-RFP surface expression and their interaction with mAb13C6 and controls. The negative control was supernatant from naïve HEK293T cultures. The positive control was a commercially available mAb13C6. Scale bar: 200 μm.

We conducted additional *in vitro* assays to evaluate the functionality of the mAb13C6 as produced in our HEK293T cultures. We constructed a plasmid (referred to here as CMV-GP-RFP) that drives the expression of a GP protein from EBOV fused with red fluorescent protein (RFP). Using this plasmid, we engineered a batch of HEK293T cells that transiently expressed GP-RFP at their cell surface (Figure 5B). Pools of these cells, expressing GP at their surface, were then used for visual evaluation of the proper binding of mAb13C6 to the GP from Ebola.

Adherent cultures of GP-RFP HEK293T cells were exposed to supernatant from cultures of mAb13C6 producers and spontaneous binding was allowed to progress for 30 min. After washing repeated times with PBS to remove unbounded mAb units, an anti-IgG conjugated with FITC was added for visual revelation of the presence of the GP-mAb13C6 complex by fluorescence microscopy (Figure 5C). We observed the expected merging of red and green, from GP and FITC, respectively (Figure 5C), in 2-D cultures of our stable clones (C66, C95, C96, and C97). These results suggest that mAb13C6, produced by our stable clones, is biologically functional, as it recognizes and specifically binds GP-EBOV displayed in cell membranes of engineered HEK293T cells.

## Concluding remarks

Preparedness to face infectious diseases is key for buffering the spreading of epidemics and minimizing the number of human deaths. Preparedness translates into simple concepts, such as accessibility of the technology and the resources to contain outbreaks efficaciously. A lesson learned from the Ebola epidemic is that we have to be fast and cost effective in our responses. This resulted in the development of new diagnostic strategies to detect EBOV in the early stages of the disease and in the introduction of efficacious therapies.

Currently, some drugs and vaccines for the treatment of EBOV are already undergoing the FDA Approval Process. Among them, mAb cocktails are recognized as one of the most promising treatments for EBOV, due to their promising results in *in vitro* experiments, in studies using animal models, and in clinical trials. However, an important consideration is that the technology needed to produce and make available sufficient doses to contain a disease outbreak has yet to be developed.

Here we presented a cost-efficient strategy that could greatly shorten the process of developing stable mammalian cell clones capable of producing monoclonal antibodies. We demonstrated the production of anti-EBOV mAbs in HEK293T cell line with activity to recognize the GP protein, in transient, stable pool, and stable clone cultures. Our transient expression cultures produced approximately 1 mg/L of mAb13C6 in a week using standard adherent cell culture techniques alone. Our further use of cost-effective selection methods that translate readily to conventional cell culture labs enabled the development of stable pools and ultimately stable clones, thereby improving the antibody production to as much as 1.8 mg/L, with specific cell productivities of 5.33 +/- 0.7 pg cell^−1^ day^−1^ in our four high-production selected clones. These specific productivity values are competitive with or higher than those previously reported for stable HEK cells, albeit still far away from the productivities needed for a commercial mAb process.^43^

Further optimization is expected from adaptation of the clones to suspended culture conditions and development of perfusion feeding protocols. These two optimization stages frequently result in a titer enhancement of at least two orders of magnitude^48^. The methodologies presented here can be directly generalized to any other mAb and easily adapted to any other glycosylated protein. In the current context of pandemic COVID-19, the urgency of the abbreviation of the process to develop stable clones is more evident than ever. ^52,53^ Our response capabilities have been cruelly challenge by COVID-19. Shortening the path for developing vaccines and neutralizing monoclonal antibodies will greatly benefit our societies.

## Acknowledgments

EGG and ARMI acknowledges funding from a doctoral scholarship provided by CONACyT (Consejo Nacional de Ciencia y Tecnología, México). GTdS and MMA acknowledge the institutional funding received from Tecnológico de Monterrey (Grant 002EICIS01). MMA, GTdS, and IMLM acknowledge funding provided by CONACyT (Consejo Nacional de Ciencia y Tecnología, México) through grants SNI 26048, SNI 256730, and SNI 1056909, respectively. The authors aknowledge the funding provided by the Federico Baur Endowed Chair in Nanotechnology (0020240I03).

